# A network modeling approach to elucidate drug resistance mechanisms and predict combinatorial drug treatments in breast cancer

**DOI:** 10.1101/176214

**Authors:** Jorge G. T. Zañudo, Réka Albert

**Affiliations:** Department of Physics, The Pennsylvania State University, University Park, Pennsylvania, 16802-6300, USA; Department of Medical Oncology, Dana-Farber Cancer Institute, Boston, Massachusetts 02215, USA; Broad Institute of Harvard and Massachusetts Institute of Technology, 7 Cambridge Center, Cambridge, Massachusetts 02142, USA; Department of Biology, The Pennsylvania State University, University Park, Pennsylvania, 16802-6300, USA

**Keywords:** breast cancer, signal transduction networks, network model, dynamic model, resistance, combination therapy

## Abstract

**Background:** Mechanistic models of within-cell signal transduction networks can explain how these networks integrate internal and external inputs to give rise to the appropriate cellular response. These models can be fruitfully used in cancer cells, whose aberrant decision-making regarding their survival or death, proliferation or quiescence can be connected to errors in the state of nodes or edges of the signal transduction network.

**Results:** Here we present a comprehensive network, and discrete dynamic model, of signal transduction in breast cancer based on the literature of ER+, HER2+, and PIK3CA-mutant breast cancers. The network model recapitulates known resistance mechanisms to PI3K inhibitors and suggests other possibilities for resistance. The model also reveals known and novel combinatorial interventions that are more effective than PI3K inhibition alone.

**Conclusions:** The use of a logic-based, discrete dynamic model enables the identification of results that are mainly due to the organization of the signaling network, and those that also depend on the kinetics of individual events. Network-based models such as this will play an increasing role in the rational design of high-order therapeutic combinations.

## I. Background

Decades of cancer research and clinical practice have showed that durable treatment of metastatic solid tumors is limited by the acquisition of resistance to the treatment (1–3). Attaining durable control of these tumors will likely require therapeutic combinations; i.e. combinations of drugs that target different key pathways within cancer cells. Our current knowledge of drug resistance mechanisms is based on resistance to single-agent treatments in cancer models and patients. The effective drug combinations employed in the clinic today, such as the ones used in chemotherapies and other notable success stories, have been mainly derived through empirical testing and following many failures. Yet the prediction of drug resistance mechanisms and design of therapeutic combinations based on scientific rationales is still an unmet need. The methods to reach these goals will have to take into account the genomic and phenotypic diversity of tumors, the variety of resistance mechanisms, and the intrinsically combinatorial nature of the problem (4–7). This makes the currently used approaches ineffective and calls for new approaches that fall within the broad umbrella of the systems biology paradigm (8,9).

Mechanistic network models of the signal transduction pathways underlying cancer cells are one of the pillars of systems biology research because of their ability to explain how these signaling cascades integrate internal and external inputs to give rise to a cellular response (10–15). These properties make mechanistic network models ideally suited to approach the problems of identifying drug resistance mechanism and designing effective hypothesis-based drug combinations. In particular, we propose using a subtype of network models, known as discrete dynamic models, which have been shown to reproduce the qualitative behavior of cancer signaling networks and are constructed solely from the regulatory interactions among the signaling proteins and the combinatorial effect of these regulatory interactions (e.g. positive or negative, additive or multiplicative) (13,16–20).

### Network models of signal transduction pathways and discrete dynamics

A signal transduction pathway consists of signaling proteins, enzymes (e.g. kinases and phosphatases), receptor proteins, and signaling molecules, which integrate extracellular and intracellular information and relay it to the transcription factors responsible for the required cellular response. Signal transduction pathways can be represented as a network, where each network node denotes an element of the signaling pathway (e.g., a signaling protein) and a directed edge between two nodes means that the first node regulates the activity of the second (target) node.

As an example, consider the simplified version of signaling through receptor tyrosine kinases (RTKs) shown in Fig. 1. In signaling through RTKs, binding of growth factors to the extracellular domain of RTKs induces a conformational change in the RTK, which promotes the recruitment and binding of several signaling proteins and kinases to its intracellular domain. Among the recruited signaling proteins are RAS and PI3K, which are activated by the RTK, and recruit other signaling molecules. RAS activates the kinase BRAF, which phosphorylates and activates the MAPK cascade (MEK/ERK), which then elicits a transcriptional response. Similarly, PI3K phosphorylates the phospholipid PIP2 into PIP3, which in turn leads to the phosphorylation of the kinase AKT, which then activates several transcription factors. Fig. 1 shows the directed interactions involved in the described sequence of events in RTK signaling, and additionally, a directed interaction between RAS and PI3K that represents the RTK-independent activation of PI3K by RAS.

**Fig. 1.**
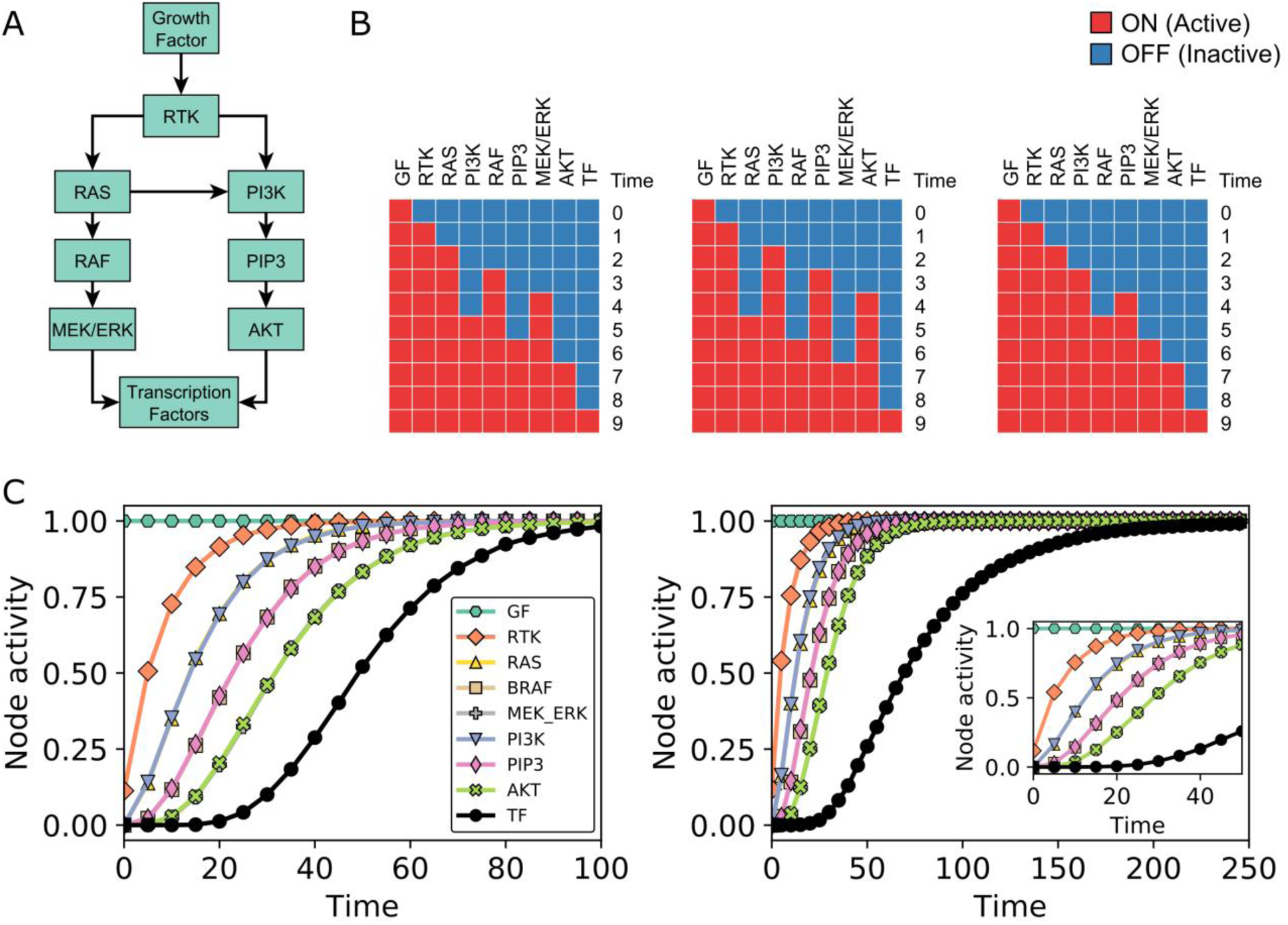
A logical dynamic network model of signal transduction. (A) Simplified network of PI3K and MAPK signaling initiated by receptor tyrosine kinases (RTKs). (B) Three possible trajectories (using general asynchronous updating) of the model constructed from the network shown in panel A and the regulatory functions in Eq. (1), with an initial condition in which the only active node is *Growth factor*(GF). (C) Time course of the activity (average node state) of each node using equal update probability for all nodes (left) or using a smaller update probability for the *Transcription factors* (TF) node compared to the rest of the nodes (right). Inset shows a zoom in of the time course for the early time steps. Note that the time courses of the activity of several nodes overlap in panels B and C, in particular, RAS and PI3K, RAF and PIP3, and MEK/ERK and AKT.

A network representation of a signaling pathway, like the one in Fig. 1, can be converted into a discrete dynamic network model by assigning a state variable σ_i_ and a regulatory function f_i_ to each node i. Each state variable σ_i_ can take a discrete number of states which denote the level of activity of the signaling element represented by node i, and where each level of activity is defined by its regulatory effect on the state variables of its target nodes. The existing experimental evidence on the number of protein conformations or post-translational modifications that yield different outcomes points to the sufficiency of assuming a small number of states, e.g. t wo or three (21,22). Each regulatory function f_i_, which can be represented using the logical operators OR, AND, and NOT, encodes the combinatorial effect on σ_i_ of the directed interactions acting on node i and thus depends on the state of the regulators of i.

As an example, we convert the network in Fig. 1A into the simplest type of discrete dynamic model, a logical (or Boolean) dynamic model, in which each node state variable σ_i_ can have two states: ON (active) or OFF (inactive). We note that the OFF state does not mean the complete absence of activity but a level of activity that is not sufficient to regulate target nodes. For the regulatory functions of this model we use the following logical rules

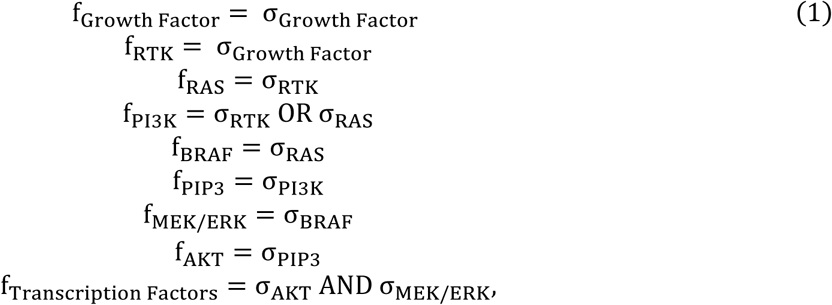

which are mathematical statements of the transmission of information between the elements in RTK signaling. For example, f_RTK_ = σ_Growth Factor_ indicates that the RTK becomes active in the presence of external growth factors, and f_PI3K_ = σ_RTK_ OR σ_RAS_ indicates that PI3K becomes active in the presence of either active RTK or active RAS. Another example is f_Transcription Factors_ = σ_AKT_ AND σ_MEK/ERK_, which indicates that the activation of the transcription factors we are considering (activation that may include transcriptional as well as post-translational regulation) requires both active AKT and active MEK/ERK. These latter rules are consistent with signaling in certain types of lung cancer (23,24).

In addition to regulatory functions like those in Eq. (1), a logical model must also specify how the node state variables change with time based on these functions, that is, we need to specify an updating scheme. Here we use the general asynchronous (GA) updating scheme (25,26,17), which updates the node state variables in discrete time units by the following two steps: (i) choosing one randomly selected node j at each time step t and updating its node state σ_j_(t) by plugging the node states of the previous time step in the regulatory function f_j_ (σ_j_(t) = f_j_[Σ(t − 1)], where Σ(t) = (σ_1_(t), σ_2_(t), …, σ_n_(t)) is the network state and encodes the state of all nodes at time t), and (ii) transferring the node state from the previous time step of the nodes not selected in step (i) (σ_i_(t) = σ_i_(t − 1), i ≠ j).

To illustrate the GA updating scheme, consider the network model in Fig. 1A, the regulatory function Eq. (1), and an initial node state Σ(t = 0) in which σ_Growth Factor_ = ON and the rest of node states are OFF. Note that the rule f_Growth Factor_ = σ_Growth Factor_ indicates that the state of the growth factor is sustained, which means that σ_Growth Factor_ will stay in its initial state (σ_Growth Factor_ = ON in this case). Three sequences of network states Σ(t) using GA updating, which we refer to as trajectories, are shown in Fig. 1B; note that, because we randomly choose one node at each time step, there are many possible trajectories. In the left-most trajectory in Fig. 1B, MAPK signaling activates before PI3K signaling, while in the right-most trajectory the activation order is reversed. The middle trajectory shows both PI3K and MAPK signaling activating concurrently. In all three cases the long-term behavior is the same: PI3K and MAPK signaling and the target transcription factors are activated, that is, all the node states in Σ are ON. Discrete dynamic models always display patterns of long-term behavior, known as dynamical attractors (e.g. steady states, such as the Σ with all nodes active of this example), which have been found to be identifiable with stable cell fates, cell states, or stable patterns of intracellular activity. The long-term behaviors (e.g. steady states) of discrete dynamic models can be identified not only by simulations, but also by alternative methods, including stable motif analysis (27,28), network reduction (29,30), and algebra-based methods (31).

The behavior of a population of cells governed by the same underlying intracellular network can be captured by the model by performing multiple simulations and interpreting each trajectory as the dynamics of a single cell. The simulated population can represent multiple types of heterogeneity by having a constitutive (in)activity of certain nodes in certain simulations, different starting states, or different kinetic parameters (this latter is implicitly captured by using stochastic update). To illustrate this latter type of heterogeneity, we use the network model of Fig. 1A and Eq. (1) with the initial condition (t=0) of Fig. 1B and perform 10,000 simulations wherein we randomly select a node with equal probability and update that node only at each time step. To capture the population-level behavior, we use the average state of node i at time t in a set of trajectories to define a quantity called the activity of the node (a_i_(t)). The activity of each node is shown in Fig. 1C left. In addition, Fig. 1C right shows how the node activity changes when incorporating the biological constraint that signaling events are faster than transcriptional events, which we do by making the probability of choosing the Transcription Factors node be smaller than that of the rest of nodes. We choose the probability to be 5 times smaller for illustration purposes, even though the difference in time scales is significantly larger; ~10^−3^-1 s for signaling events and ~10^1^-10^2^ mins for transcriptional and translational events (32,33). Both timecourses in Fig. 1C show how the network elements downstream of GF are activated sequentially in the cell population (RTK first, followed by PI3K and RAS, followed by the elements downstream of them), and how PI3K signaling and MAPK signaling are activated at the same time, on average, in the cell population. The activity of the outcome node of this simple network, the node Transcription Factors, lags behind the activity of its two regulators, as it can only activate when both AKT and MEK/ERK are active. The assumed slower timescale (lower update probability) assumed in Fig. 1C further adds to the delay of the activation of Transcription Factors (TF).

A network model can also be used to simulate the effect of drug inhibition and resistance mechanisms. For example, the addition of an RTK inhibitor in the model of Fig. 1A can be simulated by adding a node to the network denoting the RTK inhibitor (RTKi), setting the logical rule of RTK to f_RTK_ = σ_Growth Factor_ and not RTKi, and setting the state of the inhibitor to σ_RTKi_ = ON (either initially or at a certain time). Adding RTKi at time=20 causes the reversal of the increase in the activity of both branches of signaling cascades (Fig. 2B). Ultimately, all the nodes downstream of the RTK become inactive in all the simulated cells, yielding a steady state identical to the initial state (t=0 in Fig. 2B). Resistance mechanisms can be introduced in the model by changing the logical rule of the node responsible for the observed resistance, e.g., an activating RAS mutation can be introduced by setting f_RAS_ = ON. As shown on Fig. 2C, the activating RAS mutation leads to the reactivation of both signaling pathways despite the continued presence of the RTKi, and yields a steady state that differs from the steady state of Fig. 1B in the state of RTK. In other words, activating RAS mutation causes resistance to RTK inhibitors in the model. The model can be used to identify the inhibitor combinations that are able to overcome resistance. For example, the introduction of a MEK inhibitor (MEKi) at time 20 stops the continued activation of the outcome node TF (after a time delay) and leads to its inactivation in all the simulations (Fig. 2D). Thus, although the PI3K pathway is still active under this condition, from the point of view of the outcome node Transcription Factors, the combined application of RTKi and MEKi has overcome the resistance.

**Fig. 2.**
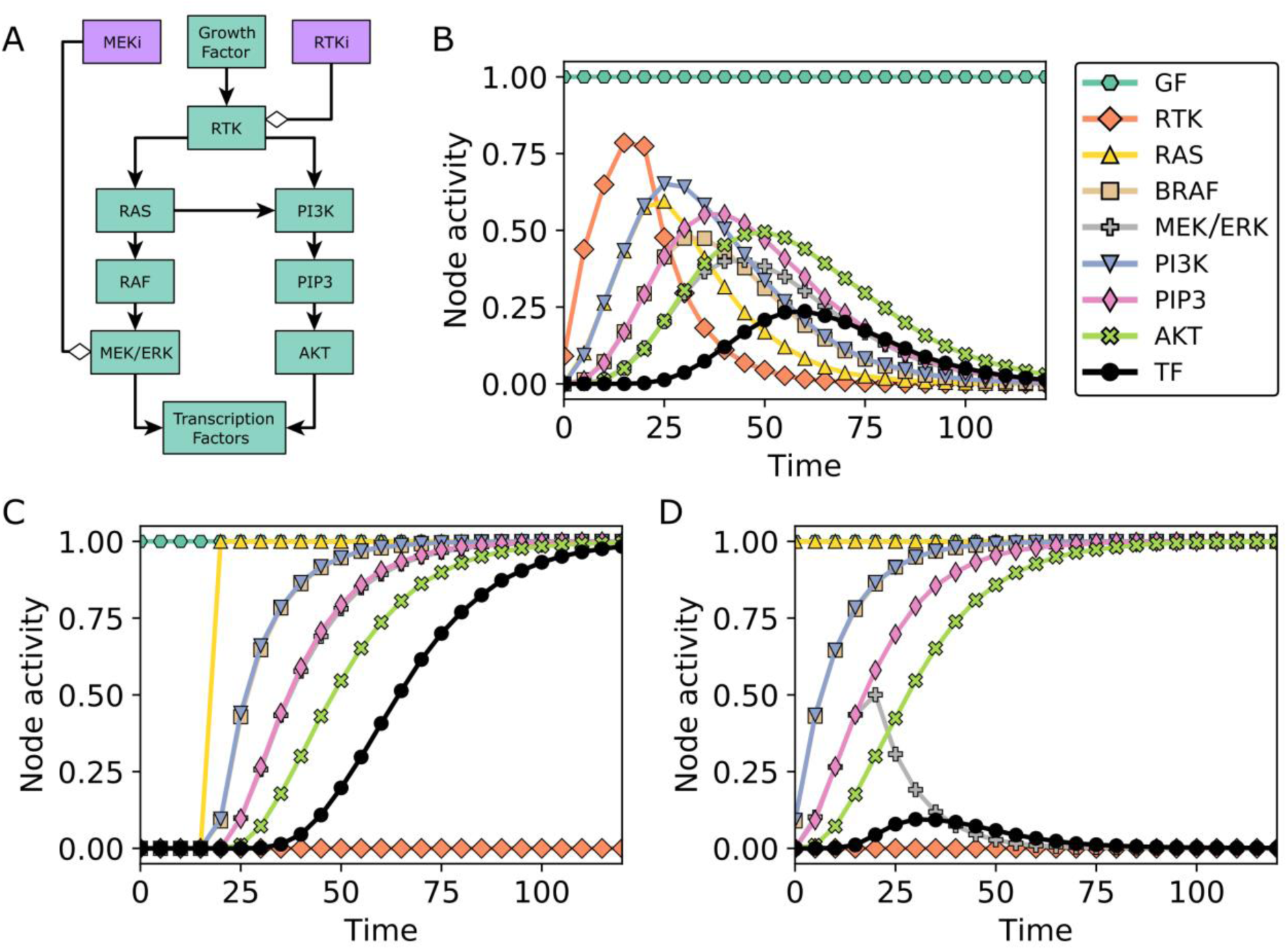
Drug inhibition and resistance mechanisms in dynamic network models. (A) The network model of Fig. 1A with additional nodes denoting a RTK inhibitor (RTKi) and a MEK inhibitor (MEKi). (B-D) Time courses of node activity (average node state) in response to Growth Factor (GF) in the presence of RTKi, a RAS activating mutation, and MEKi. For panel B, we start with an initial state in which the only active node is Growth Factor and there is no RTK inhibitor (σ_RTKi_ = OFF); we introduce the RTKi by setting σ_RTKi_ = ON for time ≥ 20. For panel C, we start with an initial state with σ_GF_ = σ_RTKi_ = ON and introduce a RAS activating mutation by setting f_RAS_ = ON for time ≥ 20. For panel D, we start with an initial state with σ_GF_ = σ_RTKi_ = ON and with the modified function f_RAS_ = ON, and introduce a MEK inhibitor by setting σ_MEKi_ = ON for time ≥ 20. Note that the time courses of the activity of several nodes overlap in panels C and D, in particular, PI3K and RAF, and PIP3 and MEK/ERK.

### Resistance mechanisms to PI3K inhibitors in breast cancer

The PI3K/AKT/mTOR signaling pathway is one of the most important regulatory pathways of cell growth and survival in healthy and cancerous cells, as evidenced by the finding that alterations in this pathway are one of the most common in human cancers (34,35). In particular, PI3KCA (an isoform of PI3K) is the most common altered gene in this pathway (mutated in ~15% of human cancers and having copy number amplifications in ~5% (35,36)) and is particularly important in the context of breast cancer (mutated in ~35% and copy number amplifications in ~5% (37–40)). The importance of PI3K in cancer has led to the development of drugs that target it. A variety of targeted drugs against PI3K are currently in clinical trials in breast cancer (34); they range in specificity from dual PI3K/mTOR inhibitors, pan-PI3K inhibitors, to isoform specific inhibitors of PI3K (e.g. Alpelisib or BYL719, a p110-alpha/PIK3CA specific inhibitor).

As a result of the development of PI3K inhibitors, there has been an increased interest in investigating the resistance mechanisms to PI3K inhibitors in the context of breast cancer, and several studies have been done in this direction (41–49). These studies have elucidated several resistance mechanisms to PI3KCA inhibitors such as PIK3CB signaling (an alternative PI3K isoform), HER3 (ERBB3) receptor activity (which is upstream of PI3K, and strongly activates the MAPK and PI3K pathway), mTORC1 signaling (which would otherwise be activated by the PI3K pathway), estrogen receptor (ER) transcriptional regulatory activity (which provides PI3K-independent means of promoting proliferation),, and signaling through the PIM (PIM1, PIM2, and PIM3), SGK (SGK1, SGK2, and SGK3), and PDK1 protein kinases (which act independently of PI3K, and have functions similar to AKT). Importantly, these resistance mechanisms have been found to be dependent on each other in some cases. For example, evidence suggests that mTORC1 signaling in BYL719 resistant breast cancer cell lines HCC1954 and JIMT1 is a consequence of the higher activity of SGK and PDK1 in these cell lines, which is sufficient to activate mTORC1 through the phosphorylation of TSC2 by SGK. This dependence between resistance mechanisms suggests that an integrative approach that fully elucidates their joint and separate mechanism of action on cell signaling and cell survival is needed for a complete understanding and to make predictions of drug interventions that overcome the observed resistance mechanisms.

## II. Results

### A network model of oncogenic signal transduction in ER+ breast cancer

We constructed a comprehensive discrete dynamic network model of signal transduction in breast cancer based on the literature of ER+, HER2+, and PIK3CA-mutant breast cancers (Fig. 3). In particular, the model incorporates the findings of resistance studies in the context of PI3K inhibitors, AKT inhibitors, RTK inhibitors, and ER inhibitors /degraders, the feedback mechanisms identified during these studies, and also includes the recent results of unbiased genome-wide screens for resistance mechanisms to PI3K inhibitors (41,43,45–66). The model consists of 51 nodes (34 Boolean and 16 multi-state nodes), of which 13 are nodes with no regulators (source nodes) that encode the initial transcriptional state of the cell (e.g. ER, HER2) or the state of nodes which are not regulated by other elements in the model (e.g. PIM and mTORC2). The nodes correspond to proteins, transcripts (8 nodes) as well as the biological outcomes proliferation and apoptosis (see Additional File 1). The edges correspond to transcriptional regulation, epigenetic mechanisms, post-translational and signaling processes. The model incorporates elements of the main signaling pathways involved in breast cancer: RTK signaling (e.g. IGF1R and HER2/HER3), PI3K signaling (e.g. PI3K and PTEN), MAPK signaling (e.g. RAS and MAPK), AKT signaling (e.g. AKT, PDK1, and FOXO3), mTORC1 signaling (e.g. mTORC1, TSC, and S6K), and ER signaling (e.g. ESR1 and MYC). These pathways converge in the survival signaling proteins that control apoptosis (e.g. BIM, BAD, and MCL1) and proliferation (e.g. cyclins, RB, and E2F), which regulate the biological-outcome nodes Proliferation (a 4-state node) and Apoptosis (a 3-state node). In addition to the 51 nodes, we also include 7 nodes that denote inhibitors of specific targets of interest^1^: Alpelisib or BYL719 (PI3K inhibitor – p110-alpha isoform specific), Ipatasertib (AKT inhibitor), Fulvestrant (ER inhibitor – selective estrogen receptor degrader (SERD)), Palbociclib (CDK4 and CDK6 inhibitor), Everolimus or Sirolimus (mTOR inhibitor), Trametinib (MEK1 and MEK2 inhibitor), and Neratinib (HER2 and EGFR inhibitor). To our knowledge, this the first comprehensive network model of its kind in breast cancer.

**Fig. 3.**
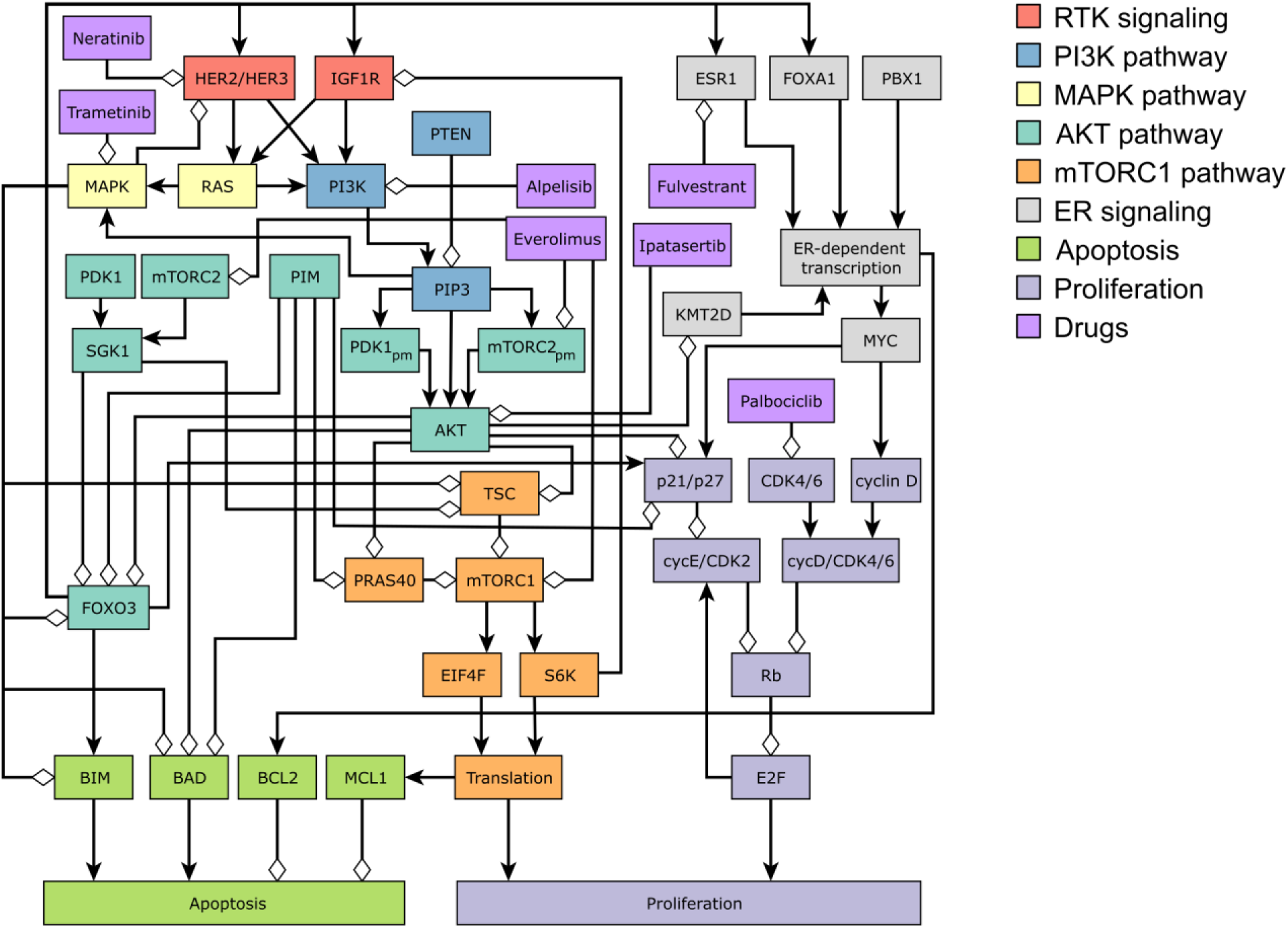
Network model of oncogenic signal transduction in ER+ breast cancer. The nodes are colored according to the pathway they are part of: RTK signaling, PI3K signaling, MAPK pathway, AKT pathway, mTORC1 pathway, ER signaling, cell-death signaling (apoptosis) and cell-cycle regulation (proliferation). The network also includes selected drugs of interest in the context of breast cancer: Alpelisib (PI3K inhibitor), Ipatasertib (AKT inhibitor), Fulvestrant (ER inhibitor), Palbociclib (CDK4/6 inhibitor), Everolimus (mTOR inhibitor), Trametinib (MEK inhibitor), and Neratinib (HER1/2 inhibitor). For clarity of the figure, we merge certain nodes into a single node when there is no ambiguity; for example, a node denoting a transcript is merged with the protein it codes, e.g., BCL2 and BCL2_T (for BCL2 transcript) are shown as BCL2. The full list of network nodes is indicated in Additional File 1.

We focus on the context of ER+/HER2-breast cancer, which we encode in the model by setting the node ER to ON and HER2 and HER3_T to OFF. In addition, we start by considering a cell state in which the source nodes IGF1R_T and PBX1 are ON (IGF1R is a common RTK in breast cancer signaling, and the subscript T denotes the intrinsic transcript level of IGF1R; PBX1 is a co-factor required for ER-dependent transcription). The source nodes PTEN, SGK1_T, PIM1, PDK1_T, and mTORC2 (which act as resistance mechanisms to PI3K inhibitors) are OFF, and the source nodes BIM_T and BCL2_T can be ON or OFF (BIM and BCL2 are pro-and anti-apoptotic proteins, respectively). In the absence of drugs, the model recapitulates a cancerous state (see Additional File 1) in which RTK, PI3K, MAPK, AKT, mTORC1, and ER signaling are active, which results in high survivability: Proliferation is high (Proliferation = 3 or 4) and Apoptosis is low (Apoptosis = 0). In this cancerous state, high survivability is possible even if the apoptotic proteins are active: anti-apoptotic protein BCL2 can be either ON or OFF, and pro-apoptotic protein BIM can be ON as long as BCL2 is also ON to counteract its effect. Thus, this state corresponds to a set of six steady states: Proliferation = 3 or 4 (caused by E2F=2 or 3), with BCL2=BIM=OFF, BCL2=BIM=ON, or BCL2=ON and BIM=OFF. This indicates high survivability states that can be either primed (BIM=ON) or unprimed for cell death (BIM=OFF), a commonly observed feature of cancer cells (67,68).

### The network model recapitulates the response to PI3K inhibitors and predicts the degree of survivability of different resistance mechanisms

We simulate the effect of a PI3K inhibitor on a population of cancer cells by starting from a combination of steady states corresponding to the cancerous state and setting Alpelisib = ON at time = 2 and maintaining it until the end of the simulation. In order to simulate the dynamics of the network model, we use general asynchronous updating, categorize nodes into fast or slow depending on whether the node is activated by a (fast) signaling event or a (slow) transcriptional/translational event, and set the update probability of fast nodes to be 5 times higher than that of slow nodes. The resulting time course of node activities is shown in Fig. 4, where time is scaled so that the time unit is equal to the average time needed to update a slow node. The time course recapitulates the experimentally observed response to PI3K inhibitors, in which PI3K inhibition has a quick and direct attenuating effect on MAPK, AKT, and mTORC1 signaling, followed by the nuclear localization of FOXO3, which transcriptionally upregulates the transcription factor ER (coded by the gene ESR1) and the pioneer factor FOXA1, which increases ER transcriptional regulatory activity. The PI3K inhibition-induced fast signal transduction events converge with the slow transcriptional events triggered by cell signaling and regulate both apoptosis and proliferation. For example, AKT and mTORC1 are quickly inhibited following PI3K inhibition, which results in the dephosphorylation and activation the pro-apoptotic protein BAD, and in the attenuation of the translational machinery. Meanwhile, pro-apoptotic protein BIM is transcriptionally up-regulated by FOXO3, and cell cycle protein cyclin D is transcriptionally upregulated due to the increased ER transcriptional regulatory activity, both of which occur later in the response to PI3K inhibition. The end result is a marked decrease in survivability: an increase in apoptosis (from Apoptosis = 0 to Apoptosis = 2 or 3, depending on whether BCL2 was initially active) and a decrease in proliferation (from Proliferation = 3 to Proliferation = 1, due to the early downregulation of MAPK, AKT, and mTORC1 activity) followed by an increase (caused by the late upregulation of ER transcriptional activity) and then stabilization at Proliferation = 2. We summarize the apoptosis and proliferation propensity with the normalized and averaged values Apoptosis_norm_ and Proliferation_norm_ (Additional File 1), which in the current simulations take the initial values Apoptosis_norm_ = 0.00 and Proliferation_norm_ = 0.50, and the final values Apoptosis_norm_ = 0.70 and Proliferation_norm_ = 0.25.

**Fig. 4.**
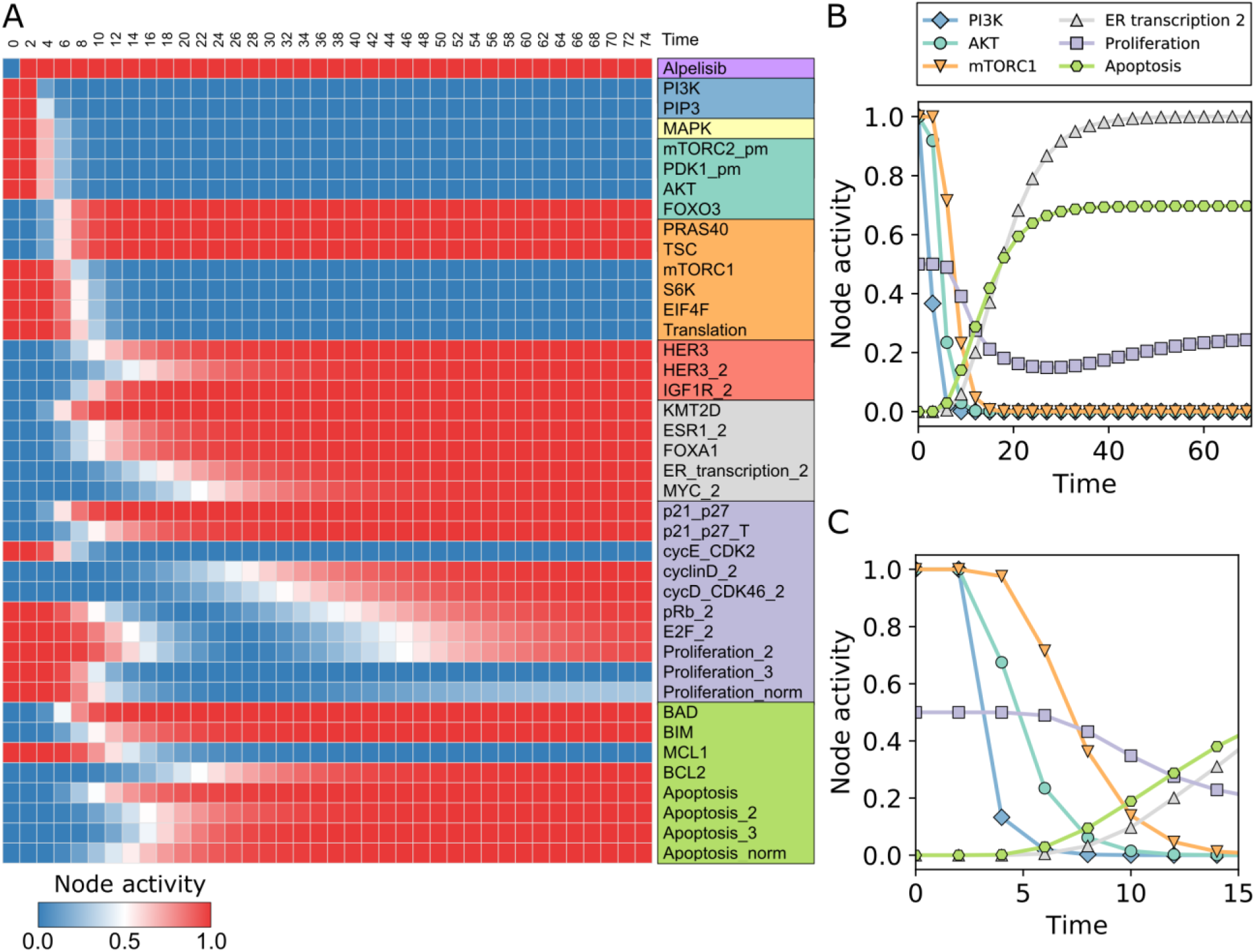
Network model response to PI3K inhibitors. (A) Time course of node activity in response to PI3K inhibition from time=2. Only the nodes that change during the time course are shown. For multi-state nodes, we show the node activity for each node state and denote each state of a multi-state node with a “_n”, where n is the state it is referring to (e.g. MAPK_1 refers to state 1 of MAPK and MYC_2 refers to state 2 of MYC). The *Apoptosis*_*norm*_ and *Proliferation*_*norm*_ is a weighted and normalized (between 0 and 1) measure of the state of the nodes *Apoptosis* and *Proliferation*. (B-C) Time course of selected nodes, each representative of a different pathway. Panel C is a zoomed in version of Panel B for the early time points.

We next test whether two recently discovered resistance mechanisms to PI3K inhibitors, PIM1/2/3 and SGK1/PDK1, increase survivability in response to PI3K inhibition in our network model. We start with an initial population of cells in the cancerous state and set either PIM = ON (which stands for any of the PIM family members) or PDK1 = SGK1_T = SGK1 = ON, and simulate the system as in the previous case (Alpelisib=ON at time=2). The resulting time course of node activities is shown in Fig. 5B-C. Both PIM and SGK1 act as resistance mechanisms to PI3K inhibitors in the model, as evidenced by a decrease in Apoptosis (from Apoptosis_norm_ = 0.70 in case of PI3K inhibitors only to 0.00/0.25 in the PIM/SGK1 cases) and an increase/lack of change in Proliferation (from Proliferation_norm_ = 0.25 to 0.50/0.25 in the PIM/SGK1 cases). A closer look at the network and the interactions of PIM and SGK1 (Fig. 5A) shows that they share most of the downstream targets of AKT, and thus, can compensate for the loss of AKT activity due to PI3K inhibition. In particular, PIM shares four out of the six AKT targets in the model (PIM does not phosphorylate TSC nor KMT2D) while SGK1 shares two AKT targets. The fact that SGK1 does not regulate the activity of pro-apoptotic protein BAD and the cyclin dependent kinase inhibitors p21/p27 is the reason why the model predicts that the PIM proteins are a stronger resistance mechanism to PI3K inhibitors compared to PDK1/SGK1. We note that this prediction depends on the relative ability of PIM and SGK1 to phosphorylate their downstream targets. To illustrate this point, Fig. 5B bottom shows the resulting time course for PIM if it is 10% less efficient than AKT on its downstream targets, (which we implement by setting PIM=OFF with a probability of 10% at every time step). While fully effective PIM maintained the cancerous mTORC1, Apoptosis_norm_ and Proliferation_norm_ values despite PI3K inhibition, in the case of the 90% effective PIM there is a decrease in the average level of mTORC1 and Proliferation_norm_ and an increase in Apoptosis_norm_, closer to the result obtained for SGK1.

**Fig. 5.**
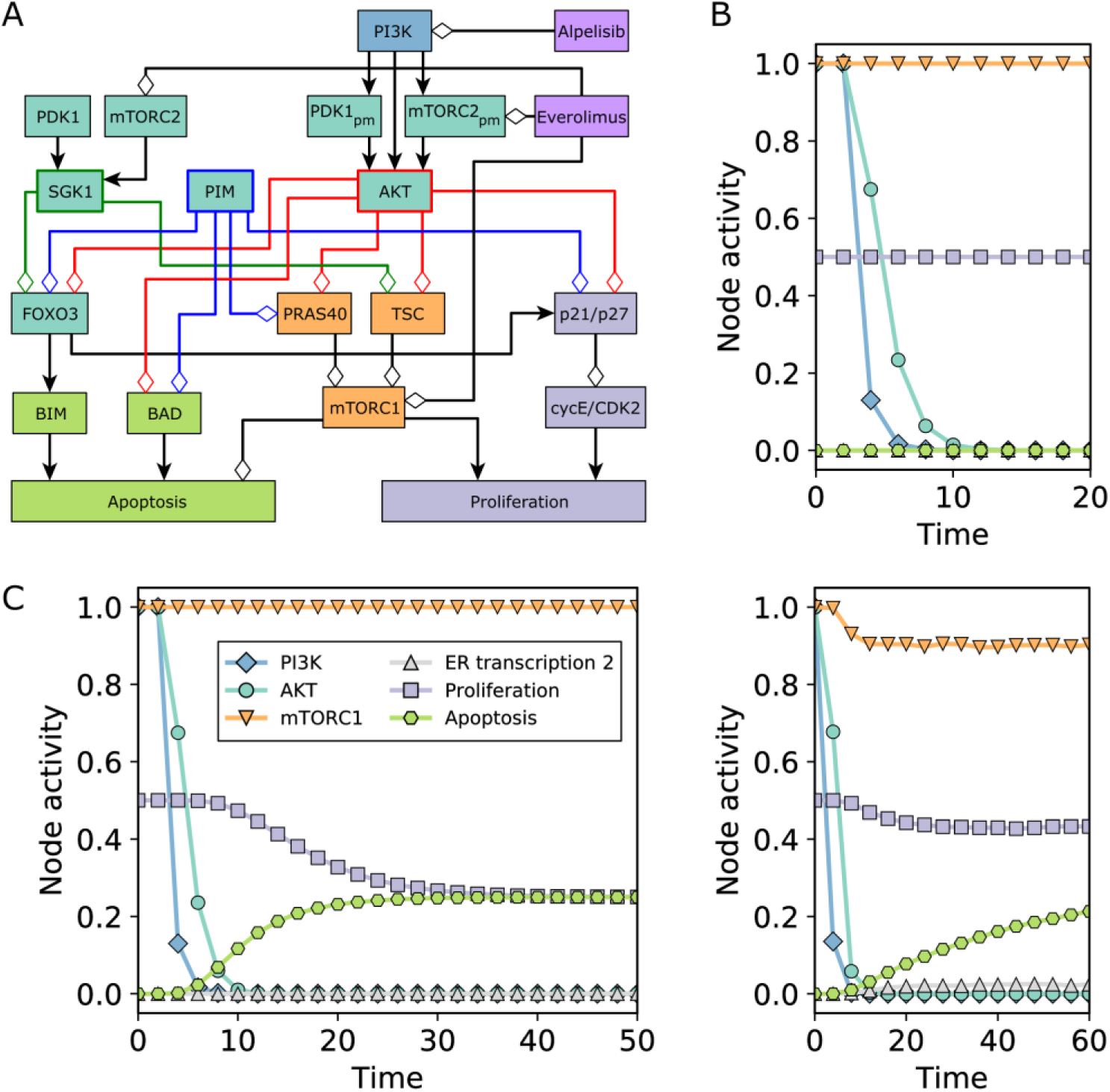
Illustration of PIM and SGK1-mediated resistance to PI3K inhibitors. (A) The relevant subnetwork of the full network shown in Fig. 3. PIM shares four targets of AKT (compare blue and red edges), while SGK1 shares two (green edges). Their post-translational regulation is different from AKT’s: SGK1 is activated by different pools of PDK1 and mTORC2 than AKT, while PIM is constitutively active. (B) Time course of node activity in response to PI3K inhibition at time=2 in cells with full activity of PIM (top) or reduced PIM activity (10% chance of inactive PIM at any time step) (bottom). (C) Time course of node activity in response to PI3K inhibition at time=2 in cells with constitutive PDK1 and SGK1 activity. The symbol legend applies to both B and C.

### The network model predicts MAPK, FOXO3, AKT, MYC, and cell cycle proteins as resistance mechanisms to PI3K inhibitors

In order to identify new resistance mechanism to PI3K inhibitors, we test every possible single and double node constitutive activation or inactivation (used in conjunction with PI3K inhibition), using an analogous procedure as in the case of PIM and SGK1/PDK1. Table 1 and Table 2 show the top node interventions that increase survivability (as measured by Apoptosis_norm_ and Proliferation_norm_) compared to the control case of PI3K inhibition with no node interventions. For the case of single node interventions, the model recapitulates the known resistance mechanisms to PI3K inhibitors: PIM, SGK1, mTORC1 (mTORC1 = ON, TSC = OFF, PRAS40 = OFF, or translation = ON), and HER2/HER3 (HER2/HER3 = 2), which lead to a decrease in the apoptosis propensity and increase in the proliferation propensity. We identify several additional resistance mechanisms: MAPK (MAPK = 1 or 2, where level 2 is the state associated with HER2/HER3 activity), which partially or fully block apoptosis, AKT (AKT = ON), which fully blocks apoptosis and restores the PI3K-inhibitor-free proliferation levels, and FOXO3 (FOXO3 = OFF or FOXO3_Ub = ON), which leads to a decrease in both the apoptosis and proliferation propensity. We also identify several resistance mechanisms that involve cell cycle proteins, namely, cyclin E and CDK 2 (cycE/CDK2 = ON), p21/p27 (p21/p27 = OFF), E2F (E2F = 3), Rb (pRb = 3), or proteins of the mitochondrial apoptosis pathway, namely BIM (BIM = OFF), BAD (BAD = OFF), MCL1 (MCL1 = ON), and BCL2 (BCL2 = ON). Several of these resistance mechanisms are supported by experimental evidence (Rb, MCL1, and BAD) or consistent with clinical observations (AKT) (46,47,49,53).

**Table 1.** Single-node resistance mechanisms to PI3K inhibitors ordered by their effect on Apoptosis. The sustained state indicated in the first column yields a decrease in *Apoptosis*_*norm*_ from 0.7 and/or an increase *Proliferation*_*norm*_ from 0.25, which are the activities of these nodes with only PI3K inhibition. Certain node perturbations that are equivalent in the network sense and lead to the same effect are grouped; specifically, PIP3=1 or 2 with PI3K=1 or 2; FOXO3_Ub=ON with FOXO3=OFF.

| Perturbation | $Apoptosis_{norm}$ | $Proliferation_{norm}$ | Mechanism |
| --- | --- | --- | --- |
| PI3K=1,2; PIP3=1,2 | 0.00 | 0.50 | PI3K |
| AKT=1 | 0.00 | 0.50 | AKT |
| HER2/HER3=2 | 0.00 | 0.50 | RTK |
| PIM=1 | 0.00 | 0.50 | AKT |
| MAPK=1 | 0.25 | 0.25 | MAPK |
| MAPK=2 | 0.00 | 0.25 | MAPK |
| BIM=0 | 0.25 | 0.25 | Apoptosis |
| BAD=0 | 0.25 | 0.25 | Apoptosis |
| SGK1=1 | 0.25 | 0.25 | AKT |
| FOXO3=0, FOXO3_Ub=1 | 0.33 | 0.13 | AKT |
| mTORC1=1 | 0.38 | 0.49 | mTORC1 |
| translation=1 | 0.38 | 0.49 | mTORC1 |
| TSC=0 | 0.38 | 0.49 | mTORC1 |
| PRAS40=0 | 0.38 | 0.49 | mTORC1 |
| MCL1=1 | 0.38 | 0.25 | Apoptosis |
| BCL2=1 | 0.50 | 0.25 | Apoptosis |
| pRb=3 | 0.69 | 0.50 | Proliferation |
| E2F=3 | 0.70 | 0.50 | Proliferation |
| cycE_CDK2=1 | 0.70 | 0.50 | Proliferation |
| p21_p27=0 | 0.70 | 0.49 | Proliferation |

**Table 2.**
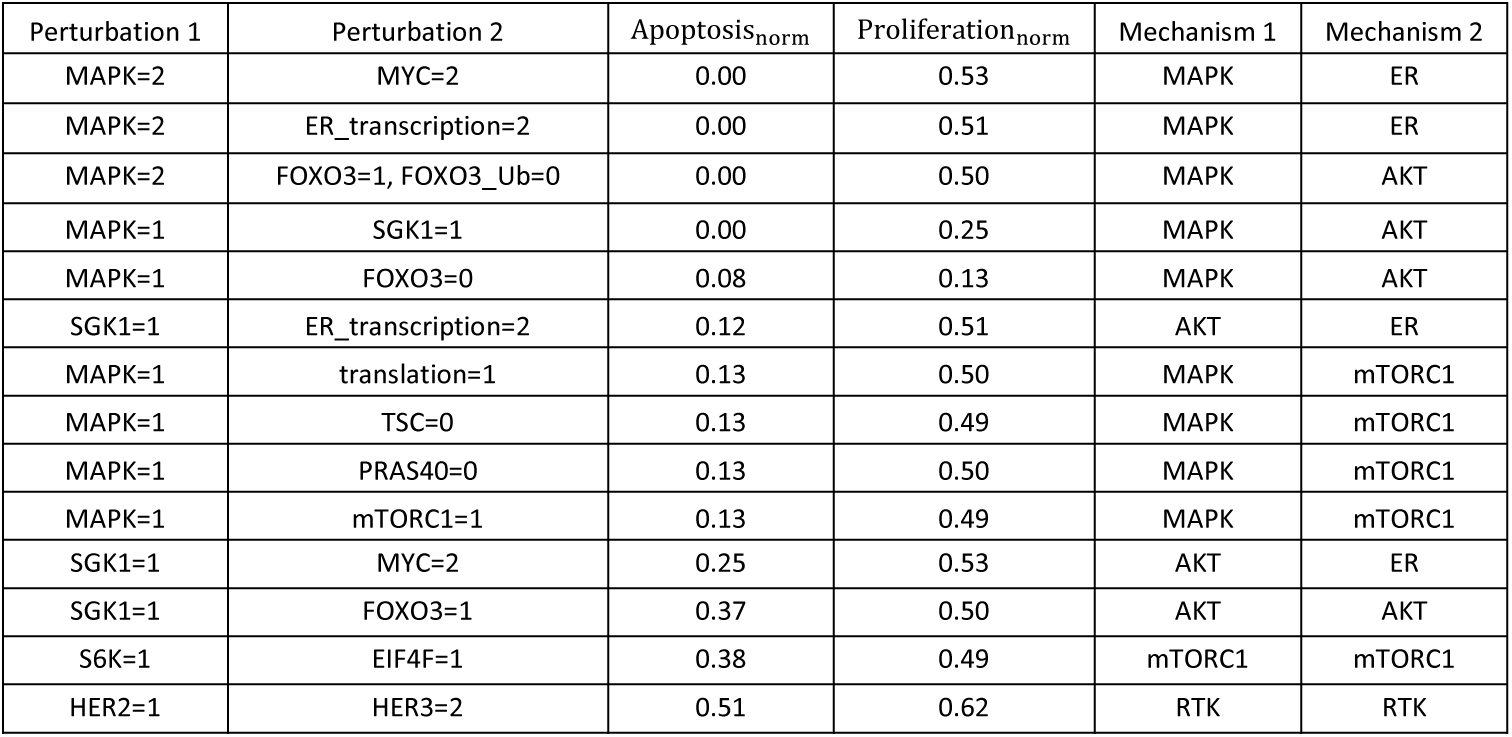
Double-node resistance mechanisms to PI3K inhibitors ordered by their effect on Apoptosis. The sustained states of the two nodes indicated in the first two columns yield a decrease in *Apoptosis*_*norm*_ from 0.7 and/or an increase in *Proliferation*_*norm*_ from 0.25 (the activities of these nodes under PI3K inhibition alone). Perturbations that involve nodes of the apoptosis or proliferation pathway are not included in this table. Certain node perturbations that are equivalent in the network sense and lead to the same effect are grouped.

For the case of double-node resistance mechanisms, and excluding those that target the apoptosis and proliferation pathways, we identify several new resistance mechanisms involving the AKT, MAPK, mTORC1, or ER pathways (Table 2). For example, MAPK = 1 combined with SGK1 = 1 fully blocks apoptosis in the model, and MAPK=2 together with high ER activity (ER_transcription = 2 or MYC = 2) restores proliferation to its PI3K-inhibitor-free level (Proliferation_norm_ = 0.50) and fully blocks apoptosis. Other examples are MAPK = 1 combined with mTORC1-activating elements (mTORC1 = 1, TSC = 0, PRAS40 = 0), which restore proliferation to its original levels (Proliferation_norm_ = 0.50) and also lower apoptosis significantly (Apoptosis_norm_ = 0.13), and FOXO3 = 1 together with MAPK = 2, which blocks apoptosis and restores proliferation (Proliferation_norm_ = 0.50).

### The network model predicts that the inhibition of the MYC-CDK4/6 axis of cell-cycle regulation and of mTORC1 synergizes with PI3K inhibitors

We next asked what single or combinatorial interventions would further sensitize cells to PI3K inhibition, i.e. yield an increased apoptosis propensity or decreased proliferation propensity compared to PI3K inhibition alone. Table 3 shows the top interventions that synergize with PI3K inhibition, which in this case are all single-node interventions. Interventions that involve the inhibition of ER activity (e.g. Fulvestrant=1, ER_transcription=0, FOXA1=0, PBX1=0, KMT2D=0) have a high anti-proliferative and apoptotic effect (Proliferation_norm_ = 0.00 − 0.13 and Apoptosis_norm_ = 0.83). Indeed, ER activity is up-regulated in response to PI3K inhibition and attenuates drug response. The synergistic effect of PI3K and ER inhibition has been previously reported (44) and is currently being explored in multiple clinical trials (34). The model predicts a novel set of combinatorial interventions that involve inhibition of PI3K and the MYC-CDK4/6 axis of cell-cycle regulation (e.g. Palbociclib=1, MYC=0, cyclinD=0, CDK4/6=0, pRb=0), which completely block proliferation (Proliferation_norm_ = 0.00) and maintain the apoptosis-inducing effect of PI3K inhibition (Apoptosis_norm_ = 0.70). Mechanistically, these novel interventions act by blocking the proliferative effect of ER, which makes their anti-proliferative effect as potent as the combination of PI3K and ER inhibitors. A second novel set of combinatorial interventions involve inhibition of PI3K and mTORC1 (e.g. Everolimus=1, mTORC1=0, S6K=0, and EIF4F=0), which is predicted to modestly increase the pro-apoptotic effect of PI3K inhibition (Apoptosis_norm_ = 0.73) and maintain its anti-proliferative effect (Proliferation_norm_ = 0.25). The synergistic effect of PI3K and mTORC1 inhibition is surprising because PI3K inhibition results in mTORC1 downregulation. We find that this combinatorial effect on apoptosis depends on the relative timing of the mTORC1 and PI3K inhibition. Early addition of mTORC1 leads to the inhibition of MCL1, which primes the cells for PI3K-inhibitor-induced apoptosis (see Table S1), and the maximum apoptosis is the same as when MCL1 is initially set to OFF.

**Table 3.** Single-node interventions which in combination with PI3K inhibitors yield an increase in *Apoptosis*_*norm*_ from 0.7 and/or a decrease in *Proliferation*_*norm*_ from 0.25. The entries are ordered by their effect on Apoptosis. Certain node perturbations that are equivalent in the network sense and lead to the same effect are grouped, specifically: ESR1=OFF, ER=OFF, and ER_transcription=0; ESR1<2 and ER_transcription<2; PBX1=OFF and FOXA1=OFF; translation=OFF, S6K=OFF, EIF4F=OFF, and mTORC1=OFF; cyclinD=0, CDK4/6=OFF, and cycD_CDK4/6=0; cyclinD<2, and cycD_CDK4/6<2; FOXO3_Ub=ON and FOXO3=OFF.

| Perturbation | $Apoptosis_{norm}$ | $Proliferation_{norm}$ | Mechanism |
| --- | --- | --- | --- |
| BCL2=0 | 1.00 | 0.25 | Apoptosis |
| ER_transcription=0 | 0.84 | 0.00 | ER |
| ER_transcription<2 | 0.84 | 0.13 | ER |
| Fulvestrant=1 | 0.83 | 0.00 | ER |
| KMT2D=0 | 0.83 | 0.13 | ER |
| FOXA1=0 | 0.83 | 0.13 | ER |
| BCL2_T=0 | 0.81 | 0.25 | Apoptosis |
| MCL1=0 | 0.75 | 0.25 | Apoptosis |
| mTORC1=0 | 0.74 | 0.25 | mTORC1 |
| Everolimus=1 | 0.73 | 0.25 | mTORC1 |
| cycD_CDK46=0 | 0.70 | 0.00 | Proliferation |
| Palbociclib=1 | 0.70 | 0.00 | Proliferation |
| pRb=0 | 0.70 | 0.00 | Proliferation |
| E2F=0 | 0.70 | 0.00 | Proliferation |
| MYC=0 | 0.70 | 0.00 | ER |
| pRb<2 | 0.70 | 0.13 | Proliferation |
| cycD_CDK46<2 | 0.70 | 0.13 | Proliferation |
| MYC<2 | 0.70 | 0.13 | ER |
| FOXO3=0 | 0.33 | 0.13 | AKT |

## III. Discussion

Network models excel in both aspects of model utility: the integration and interpretation of existing knowledge, and the generation of novel predictions. Our network model (Fig. 3) unites information from numerous studies and visualizes the inter-relationships among various pathways and processes. The overlay of the usually-defined pathways (marked by separate colors) on the signal transduction network that starts with receptor tyrosine kinases and ends with two phenotypic outcomes reveals that these pathways cover a variety of subgraphs of the full network, from linear cascades (such as RAS-MAPK) to bow-tie structured neighborhoods of a node (such as ER signaling) and to subgraphs wherein negative regulation or feedback plays an important role. The inter-regulation among subgraphs is also substantial, and biologically significant. To better illustrate this point, in Fig. 6A we represent each the seven pathways relevant to ER+, PI3K mutant breast cancer and the aggregated relationships between them. These relationships summarize one or multiple paths between the pathways. For example, mTORC1-induced protein translation, which leads to the increased activity of the anti-apoptotic protein MCL1, yields an overall negative regulation between the mTORC1 pathway (orange rectangle) and the apoptosis pathway (green rectangle). The positive effect of the AKT pathway on the mTORC1 pathway summarizes AKT and PIM’s inhibition of PRAS40, as well as AKT and SGK1’s inhibition of TSC; both PRAS40 and TSC are inhibitors of mTORC1.

**Fig. 6.**
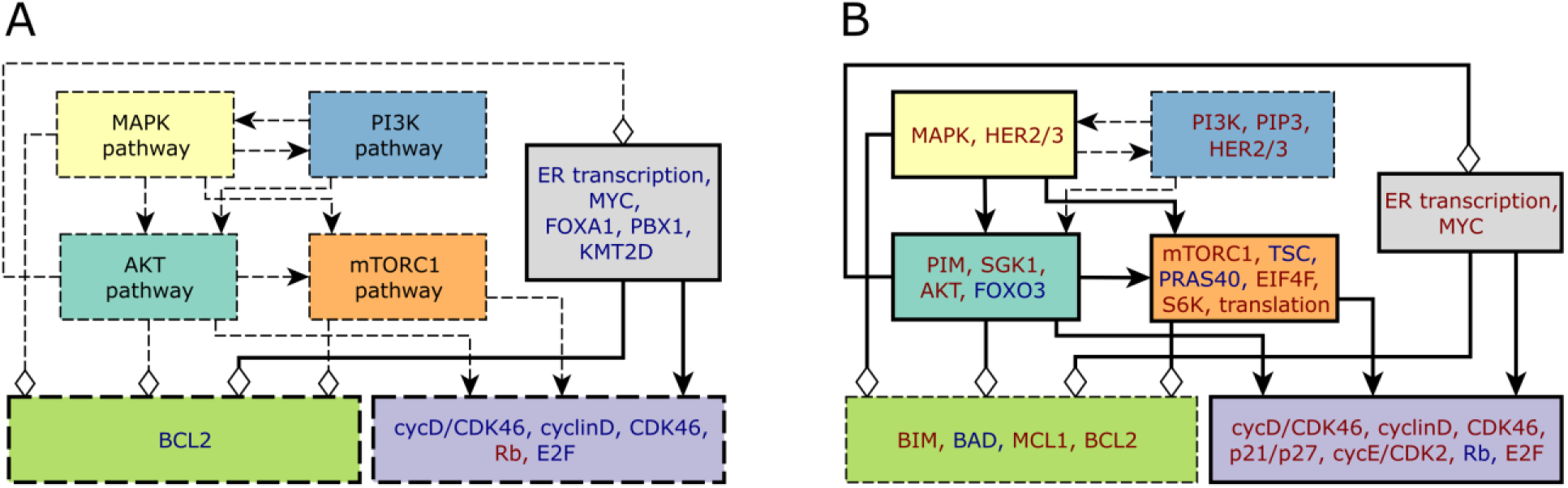
Synergistic interventions and resistance mechanisms to PI3K inhibitors. The colored rectangles correspond to the pathways introduced in Fig. 3. Thick continuous lines indicate active pathways/interactions, thick and dashed lines represent partially active pathways/interactions, thin and dashed lines mean inactive pathways/interactions. For the node names indicated inside the colored rectangles, blue indicates inhibition/inactivity and red indicates increased activity. (A) Signaling pathway activity in response to PI3K inhibition. ER signaling is still active (partly due to the release of its inhibition by AKT), while the apoptosis and proliferation pathways are partially active. Inhibition of the nodes indicated in blue font or constitutive activity of Rb is predicted to have a synergistic effect with PI3K inhibition. (B) Resistance mechanisms to PI3K inhibitors. Sustained activity of the nodes indicated with red font inside each pathway can (at least partially) restore the pathway’s activation and obstruct the effectiveness of PI3K inhibition. Sustained inactivity of the nodes indicated with blue font can have a similar effect. For simplicity, the HER2/HER3 resistance mechanism is not included in a separate RTK module but as part of the pathways activated by HER2/HER3, namely MAPK and PI3K.

AKT inhibits ER signaling through downregulating the histone methyltransferase KMTD and by its inhibition of FOXO3, which would otherwise activate ER. This negative edge stands out from and opposes an otherwise sign-consistent meta-network, wherein the five upstream pathways have positive inter-regulation and all favor proliferation and/or disfavor apoptosis. In ER+, PI3K mutant breast cancer cells, this negative edge dampens (but does not block) ER signaling, and all four other pathways are active; yielding the collective effect of a significant proliferation propensity and lack of apoptosis propensity. In case of drug inhibition of PI3K, four pathways (PI3K, MAPK, AKT, mTORC1) are inactivated, and consequently the break on ER signaling is released (Fig. 6A). The overall effect is a significant apoptosis propensity and a low (but non-zero) proliferation propensity. While the quantification of the two biological outcomes depends on specific model and implementation details, the main message is clearly encapsulated in the network: PI3K inhibition does not eliminate all the proliferation-inducing, apoptosis-resisting activity in the network. Our model provides specific predictions on what additional interventions would yield a significant improvement over PI3K inhibition alone. The targets of these predicted interventions lie in the ER signaling, cell cycle and apoptosis pathways; their names and the nature of their control (inhibition or activation) is also indicated in Fig. 6A. Our finding that multiple combinatorial interventions are effective enables the selection of those that are most effective drug targets and minimize toxicity and side effects.

A novel prediction of the model is that PI3K + CDK4/6 inhibition is a very effective combination treatment because of its ability to both induce cancer cell death and cell cycle arrest by suppressing two parallel proliferation regulator complexes (cyclin E and CDK2, and cyclin D and CDK4/6). So far there are few studies of the combined effect of PI3K inhibitors and CDK4/6 inhibitors (47,69), and the on-going clinical trials all include ER inhibitors simultaneously with both PI3K and CDK4/6 inhibitors (34,69). The model predicts that the combination of PI3K and CDK4/6 inhibitors can be as effective as the combination of PI3K and ER inhibitors, and that the addition of CDK4/6 inhibitors to the latter does not further increase its effectiveness. Given that resistance to ER inhibitors can be overcome by CDK4/6 inhibitors (66) and that targets of CDK4/6 inhibitors are known resistance mechanisms to ER inhibitors (70,71), the model predictions suggests that PI3K inhibitors + ER inhibitor followed by PI3K inhibitors + CDK4/6 inhibitors after acquisition of resistance is a better strategy than combined PI3K + ER + CDK4/6 inhibitors.

The network of inter-relationships among pathways can also be used to interpret the existing information and new predictions on potential resistance mechanisms to PI3K inhibition in PI3K mutant, ER+ breast cancer (Fig. 6B). Broadly speaking, any mechanism that yields the restoration of activity in the PI3K, MAPK, AKT or mTORC1 pathways, or increased activity of the ER pathway, will restore the proliferation-inducing and/or apoptosis-opposing effects of these pathways, and will yield a decrease in the effectiveness of PI3K inhibitors. For example, a mechanism that would partially restore PI3K activity or PIP3 levels (for example by a loss of function alterations in PTEN (72)), could lead to the restoration of the PI3K→ AKT and PI3K → MAPK edges in Fig. 6B and thus reverse the effects of PI3K inhibition. Constitutive activity of AKT, PIM or SGK1, or inhibition of FOXO3, would at least partially restore the four outgoing edges of the AKT pathway. While one of these edges is to dampen ER signaling, the other three will lead to a decreased apoptosis propensity and increased proliferation propensity. Inspecting the multitude of potential resistance mechanisms (indicated by color-coded node names inside each pathway symbol in Fig. 6B), those in the PI3K, AKT and mTORC1 pathways may be categorized as pathway reactivation, if we consider the union of these three, i.e. PI3K/AKT/mTORC1, as the index pathway. Constitutive ER transcriptional regulatory activity is an example of pathway bypass: it leads to cell cycle progression and activates the anti-apoptotic protein BCL2. Constitutive activity of the MAPK pathway, a model-predicted resistance mechanism, resembles pathway bypass in that it inhibits FOXO3, which would otherwise be accomplished by AKT, but it also overlaps the index pathway through its activation of mTORC1. The model predicts that two different combinations of MAPK and FOXO3 activity (FOXO3=ON and MAPK=2 or FOXO3=OFF and MAPK=1) can both act as resistance mechanisms to PI3K inhibitors. An analysis of the elements regulated by MAPK and FOXO3 reveals that this happens because two normally opposing effects are allowed to co-occur. In the FOXO3=ON and MAPK=2 case, MAPK’s regular inhibition of FOXO3 is blocked, thus this combination yields the proliferative effect of FOXO3=1 but a lesser pro-apoptotic effect (due to MAPK=2). In the FOXO3=OFF, MAPK=1 case the apoptosis propensity is decreased because MAPK=1 inhibits BAD.

Here we focused on PI3KCA mutant breast cancer and targeted PI3K inhibition, which is showing promising results in clinical trials. Our network modeling framework can be used or adapted to answer a broader set of questions. For example, we can consider mutants that have one of the model-identified resistance mechanisms, determine the drugs that overcome the resistance, and identify the most effective combinatorial therapies. An example of such a prediction is that patients with activating genetic alterations in PIM would greatly benefit from the combination of PI3K inhibitors with PIM inhibitors (as one would expect) or the combination of ER and mTOR inhibitors. Although we focused on ER+/HER2-breast cancer, the pathways and mechanisms of the HER2+ subtype are included in the model. Indeed, HER2/HER3 appears as a resistance mechanism to PI3K inhibition and the model recapitulates several resistance mechanisms observed in HER2+ breast cancer (Additional File 1). The model can be expanded to incorporate additional resistance mechanisms relevant to HER2+ breast cancer (e.g. the FGFR signaling pathway in the context of estrogen receptor degraders (73–75)).

Certain predictions of the model rely on considerations that go beyond the network structure, for example timing. The model predicts a non-monotonic decrease in proliferation in response to PI3K inhibition (Fig. 7). This is due to the convergence of fast signal transduction events that decrease proliferation with the slow ER-driven transcriptional events that increase it. Timing also plays a key role in the predicted synergistic effect on apoptosis induction of mTORC1 inhibition followed by PI3K inhibition. This is because mTORC1 inhibition leads to the inhibition of MCL1, which primes the cells for PI3K-inhibitor-induced apoptosis (see Table S1). The observation that the timing of drugs can prime cells for apoptosis and yield drug synergy is consistent with previous work showing a similar effect in triple-negative breast cancer (68).

## IV. Conclusions

The breast cancer network model we present in this work integrates the current knowledge of PIK3CA-mutant, ER+ and HER2+ breast cancers, and uses it to identify a set of elements that may eventually be exploited in high-order therapeutic combinations to achieve a more durable control of breast cancer. The breast cancer model and its predictions will serve as a basis for guiding and interpreting drug resistance and drug combination studies in the context of ER+ and/or HER2+ breast cancer. The model can be expanded to incorporate multiple genetic alterations observed in breast cancer patient cohorts (38,40,76,77) by introducing these alterations into the model appropriately (e.g., as a node activation or inactivation). The inclusion of the most probable intrinsic or acquired resistance mechanisms to a treatment, informed by pre-, on-and post-treatment genetic characterization of tumors (77), will allow the identification and ranking of the combinatorial interventions that are effective even in the presence of tumor drug resistance. In the future, we expect experimentally and clinically validated network models similar to the one presented here to be an integral part of precision medicine, and to be able to identify successful combinatorial therapies in predicted cell subpopulations and in tumor types and subtypes of interest.

## V. Methods

### Model simulations

The simulations of the discrete network models were done using the BooleanDynamicModeling Java library, which is freely available on GitHub (https://github.com/jgtz/BooleanDynamicModeling). To simulate multi-level nodes, we use a Boolean variable to denote each level greater than 1. For example, for a 3-level node with states 0, 1, and 2, we would have 2 variables (Node and Node_2), and for a 4-level node we would have 3 variables (Node, Node_2, and Node_3). The regulatory functions of all the nodes are indicated and explained in Additional File 1.

### Attractor-finding in discrete network models

To attractors of the model were identified using the StableMotifs Java library, which is freely available on Github (https://github.com/jgtz/StableMotifs) and implements the attractor-finding method based on stable motif analysis (27,28), as has been previously described (17). Stable motif analysis can find the attractors of a logical model by identifying the model’s stable motifs, a set of nodes and their node states with certain identifiable topological (intersecting directed cycles) and dynamical properties (partial steady states), which uniquely determine the attractors of the model (27,28).

## List of abbreviations

AKT: protein kinase B
ER: estrogen receptor
HER2: human epidermal growth factor receptor 2;
MEK: mitogen-activated protein kinase kinase
MEKi: MEK inhibitor
PI3K: phosphatidylinositol – 4,5-biphosphate 3-kinase
RTK: receptor tyrosine kinase
RTKi: RTK inhibitor

The full names of the abbreviated node names in Fig. 3 and thereafter are indicated in Additional File 1.

## Ethics approval and consent to participate

Not applicable

## Consent for publication

Not applicable

## Availability of data and materials

All the data are contained in the text and additional files.

## Authors’ contributions

J.G.T.Z. and R.A. designed research; J.G.T.Z. and R.A. designed the model; J.G.T.Z. performed the analyses; and J.G.T.Z. and R.A. wrote the paper.

## Acknowledgements

J.G.T.Z. would like to thank Levi Garraway and Nikhil Wagle for hosting him as a visiting Research Fellow at the Dana-Farber Cancer Institute and Broad Institute of Harvard and MIT. We would also like to thank Maurizio Scaltriti and Pingping Mao for their helpful suggestions on the manuscript. This project is part of a group effort by the Stand Up to Cancer Drug Combination Convergence Team; we thank the members of the team for constructive discussions and feedback.

## Additional Files

**Table S1.**
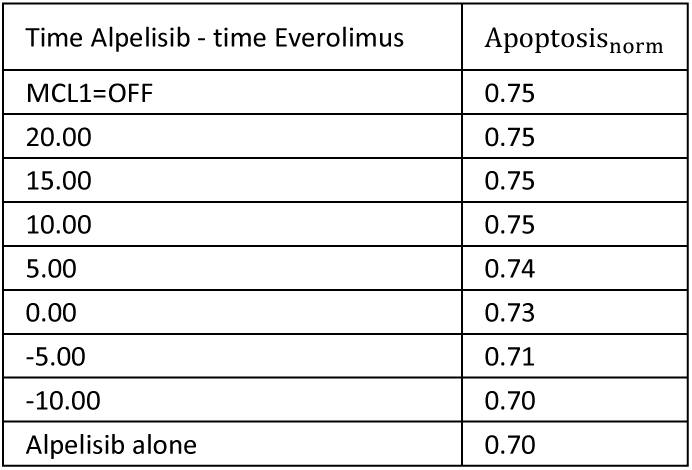
Timing-dependent combinatorial effect of PI3K inhibitors (Alpelisib) and mTOR inhibitors (Everolimus). We demonstrate that the combinatorial effect on apoptosis depends on the relative timing of the mTOR and PI3K inhibition by doing simulations in which we add the mTOR inhibitor at varying times before the PI3K inhibitor.

### Additional File 1

#### Explanation of the node names

ESR1 – the transcript of the *ESR1* gene, which encodes the estrogen receptor

ER-estrogen receptor status of the cell

HER2-human epidermal growth factor receptor 2 status,

HER3-human epidermal growth factor receptor 3,

HER2/HER3 – heterodimer of HER2 and HER3

IGF1R – insulin-like growth factor 1 receptor (IGF1R) and insulin receptor (INSR)

IGF1R_T – transcript of IGF1R or INSR

PIP3 – phosphatidylinositol – (3,4,5)-triphosphate

PI3K – phosphatidylinositol – 4,5-biphosphate 3-kinase, encoded by the gene *PIK3CA*

PTEN – phosphatase and tensin homolog

AKT – protein kinase B

RAS – Ras family small GTPase,

MAPK – merged node that includes RAF-rapidly accelerated fibrosarcoma kinase family,

MEK-mitogen-activated protein kinase kinase, and ERK-extracellular signal-regulated kinase

FOXA1 – forkhead box protein A1

PBX1 – pre-B cell leukemia transcription factor 1

KMT2D – histone-lysine N methyltransferase 2D

MYC-transcription factor encoded by the proto-oncogene *c-myc*

PDK1 – 3-phosphoinositide dependent protein kinase 1, encoded by the gene *PDPK1*

PDK1_pm-3-phosphoinositide dependent protein kinase 1 localized in the plasma membrane

SGK1 – serum and glucocorticoid-induced kinase 1

SGK1_T – transcript of serum and glucocorticoid-induced kinase 1

PIM – Pim-1 Proto-Oncogene, Serine/Threonine Kinase (PIM1), Pim-2 Proto-Oncogene, Serine/Threonine

Kinase (PIM2), and Pim-3 Proto-Oncogene, Serine/Threonine Kinase (PIM3).

cycD/CDK4/6 – complex formed by cyclin D1 and cyclin dependent kinase 4/6,

cycE/CDK2 – complex formed by cyclin E and cyclin dependent kinase 2,

Rb – retinoblastoma family proteins

E2F-transcriptional activator members of the E2F family

p21 – WAF1/CIP1, cyclin-dependent kinase inhibitor 1, encoded by the gene *CDKN1A*

p27-KIP1, cyclin-dependent kinase inhibitor 1B, encoded by the gene *CDKN1B*

mTORC1-mechanistic target of Rapamycin complex 1

mTORC2_pm-mechanistic target of Rapamycin complex 2 localized in the plasma membrane

mTORC2-mechanistic target of Rapamycin complex 2 not localized in the plasma membrane and possibly localized in the mitochondria.

TSC – tuberous sclerosis complex 1 (TSC1) and tuberous sclerosis complex 2 (TSC2)

PRAS40-proline-rich Akt substrate of 40 kDa, a component of mTORC1

FOXO3-forkhead box O3 protein

FOXO3_Ub – forkhead box O3 protein degraded through the ubiquitin proteasome pathway

S6K – p70 ribosomal S6 kinase

EIF4F-eukaryotic initiation factor 4A(EIF4A), eukaryotic initiation factor 4G (EIF4G), and Eukaryotic

Translation Initiation Factor 4E Binding Protein 1 (4EBP1)

Translation – processes related to ribosome translation, cap-dependent translation

Proliferation – focusing on two requirements: cell cycle transition from G1 to S, and protein translation

BCL2-B-cell lymphoma 2 protein, anti-apoptotic family representative

BCL2_T – B cell lymphoma 2 transcript

MCL1 – BCL2 family apoptotic regulator, anti-apoptotic

BAD-Bcl-2-associated death promoter, apoptosis sensitizer family representative

BIM-Bcl-2-like protein 11, apoptosis activator family representative

BIM_T – transcript of BCl-2-like protein 11

Alpelisib – Isoform-specific drug inhibitor of PI3K alpha, also known as BYL719

Fulvestrant – drug inhibitor of the estrogen receptor, a type of selective estrogen receptor degrader (SERD)

Neratinib – drug inhibitor of EGFR and HER2

Palbociclib – drug inhibitor of CDK4/6

Everolimus and Everolimus analogues (Temsirolimus, Everolimus) – mTORC1 and mTORC2 inhibitors

Trametinib – drug inhibitor of MAPK signaling (specifically, of MEK1 and MEK2)

Ipatasertib – drug inhibitor of AKT

#### Explanation of the node states

To maximize the parsimony of the network, we aim to represent each gene product with the minimal number of nodes. This can be a single node if there is a single main mode of regulation of that gene (e.g. transcriptional). If there is both transcriptional and (post)translational regulation, we have a node for the transcript (marked by the subscript “T”) and a separate node for the protein. In the case of PDK1 and mTORC2 there is evidence of two different pools (localizations) that are regulated differently (i.e. the plasma membrane localized PDK1 and mTORC2 are regulated by PIP3) and have different effects (i.e. one localization is mainly responsible for the regulation of AKT, while the other is mainly responsible for the regulation of SGK1). For this reason, we define two nodes for PDK1 and mTORC2. Finally, the post-translational modifications of FOXO3 can be categorized into faster processes (phosphorylation-induced translocation) and the slower process of phosphorylation-induced proteolytic degradation. Because of this reason we define a separate node, FOXO3_Ub, and update it with a lower probability.

For the 34 binary nodes, the node name implicitly represents the active state of the node. For example, RAS refers to the GTB-bound active form of the RAS GTP-ase. If a protein is activated by phosphorylation (such as AKT, S6K, SGK1), the node name refers to the phosphorylated form, and the regulatory function expresses the condition under which the protein is activated (e.g. AKT is phosphorylated and activated by PDK1 and mTORC2). If a protein is active if unphosphorylated (such as p21, p27, FOXO3, BAD, BAX), the node name refers to the unphosphorylated form; the network and the regulatory function will reflect phosphorylation as a negative regulation of this node.

The node states of the receptors ER and HER2 reflect the receptor expression status of the cell; i.e. an ER+ cell would have ER=1.

Multi-level nodes characterize the receptor tyrosine kinases IGFR1, HER3, the HER2/HER3 heterodimers, the PI3K pathway, RAS and the MAPK pathway, the ER pathway, the majority of the cell cycle related proteins (e.g. cyclin D, Rb, E2F), and the outcome nodes proliferation and apoptosis. For these 16 multi-level nodes, separate regulatory functions are given for the basal activity, activity level 2 (denoted by “_2”), and higher activity levels if they exist. The regulatory functions of the higher activity levels reflect a more stringent condition; thus, when a node is activated at a level i>1, it is also activated at level i-1.

In the case of Rb, which is active in the unphosphorylated form, we introduce a multi-level node pRb to denote the (hyper)phosphorylated form of Rb.

The output nodes “Proliferation” and “Apoptosis” reflect the propensity of a cell to commit to cell cycle progression, and programmed cell death, respectively. The higher the node level, the stronger this propensity is. In order to facilitate comparison, we introduce normalized variables “Proliferation_norm_” and “Apoptosis_norm_”. The normalization is exponentially weighted following the formula *x*_*norm*_(*x* = *i* = 1/2^*n-i*^, where i=1,… n, n is the maximal activity level of node *x*, and *x*_*norm*_(*x* = 0. When performing multiple simulations, we report the average of the normalized proliferation and apoptosis values over the ensemble of simulations. These values can be loosely interpreted as estimates of the fraction of a simulated cell population that is undergoing cell cycle progression, or is on the path toward apoptosis, respectively. Rather than interpreting the values themselves, we focus on the sign of the change (i.e. increase or decrease) in these values following an intervention or perturbation.

#### Regulatory functions in the model

The 10 nodes that are not regulated by other network nodes or by drugs (BCL2_T, BIM_T, ER, IGF1R_T, HER3_T, PBX1, PDK1, PIM, PTEN, SGK1_T) are assumed to maintain their initial state. The regulatory function of these nodes is of the type *f*_*X*_ = σ_*X*_, meaning that the future state of node X equals its current state. We do not explicitly indicate these regulatory functions. We do indicate the regulator functions of the three source nodes that are regulated by drugs (CDK4/6, HER2, mTORC2).

In the regulatory functions that follow for simplicity we represent the node state by the node name (i.e. we use X instead of σ_X_).

f_IGF1R_=IGF1R_T or (HER2 and FOXO3)

f_IGF1R_2_=(IGF1R_T or (HER2 and FOXO3)) and not S6K

A basal activity of IGF1R is achievable due to an intrinsic transcript level or by transcriptional upregulation by FOXO3 (PMID: 21215704). The transcriptional upregulation of IGFR1 by FOXO3 depends on the presence of HER2, since it has been found in HER2^+^ cells (PMID: 21215704). In order to activate IGF1R at a higher level S6K must be absent, as S6K indirectly inhibits IGF1R through IRS1 (PMID: 16452206).

f_HER3_T_= HER3_T

f_HER3_=FOXO3 or HER3_T

f_HER3_2_=FOXO3

FOXO3 transcriptionally activates HER3 above its intrinsic level (PMID: 21215704).

f_mTORC2_=mTORC2 and not Everolimus

mTORC2 maintains its activity unless inhibited by Everolimus analogues.

f_SGK1_=SGK1_T and PDK1 and mTORC2

Active SGK1 protein needs to be translated from its transcript and needs to be double phosphorylated by PDK1 and mTORC2 (PMID: 20027184, PMID: 18925875).

f_HER2_= HER2 and not Neratinib

f_HER2/HER3_= HER2 and (HER3 or HER3_2) and not Neratinib

f_HER2/HER3_2_= HER2 and ((HER3 and not MAPK_2) or HER3_2) and not Neratinib

HER2 forms heterodimers with HER3 (PMID: 11252954), unless inhibited by Neratinib. Strong activation of MEK inhibits the activity of the heterodimer (PMID: 22552284). We assume that high activation of HER3 is able to overcome the inhibition by MAPK. This is supported by the observation that in HER2+ breast cancer AKT inhibition leads to an increase in MAPK activity (PMID:24436048). This increase is mediated by the activation of FOXO3, which transcriptionally upregulates HER3. For the increased MAPK activity to be compatible with HER3 and HER2/3 heterodimer activity, a sufficiently high level of HER3 must be able to overcome MAPK inhibition.

f_RAS_=IGF1R or IGF1R_2 or HER2/HER3 or HER2/HER3_2

f_RAS_2_=HER2/HER3 or HER2/HER3_2

f_RAS_3_=HER2/HER3_2

IGF1R and HER2/HER3 activate RAS (PMID: 17496910, PMID: 21531565).

f_MAPK_=(RAS or RAS_2 or RAS_3) and (PIP3 or PIP3_2) and (not Trametinib or RAS_3)

f_MAPK_2_=(RAS_2 or RAS_3) and PIP3 and (not Trametinib or RAS_3)

RAS activates RAF by translocating it to the plasma membrane, where protein kinases phosphorylate and activate RAF (PMID: 17496910). RAF activates MEK by phosphorylation (PMID: 8325833, PMID: 17496910). PIP3 can indirectly activate MEK (PMID 24327733); the importance of this mechanism (and the use of the “and” operator) is supported by the fact that PI3K inhibition leads to rapid downregulation of ERK phosphorylation (PMID: 24436048). ERK is activated by double phosphorylation by MEK (PMID: 2032290). Higher activity of RAS yields higher activation of the MAPK pathway, and the highest RAS activity can overcome the inhibitory effect of Trametinib. This rule is consistent with the observation that drug inhibition of MEK in HER2+ cells leads to increased HER2/HER3 heterodimer formation and to reactivation of the MAPK pathway 24 hours after drug treatment (PMID: 22552284, PMID: 24436048).

f_PI3K_=(IGF1R or IGF1R_2 or HER2/HER3 or HER2/HER3_2 or RAS or RAS_2 or RAS_3) and (not Alpelisib or HER2/HER3_2)

f_PI3K_2_= HER2/HER3_2 and not Alpelisib

PI3K is activated by IGF1R, the HER2/HER3 heterodimer or RAS (PMID: 16452206, PMID: 11252954, PMID: 12853564, PMID: 8052307). It is inhibited by the drug Alpelisib (also known as BYL719). We assume that high activation of HER2/HER3 can overcome the inhibitory effect of Alpelisib, and reactivate PI3K.

f_PIP3_=(PI3K or PI3K_2) and not PTEN

f_PIP3_2_=PI3K_2 and not PTEN

PI3K catalyzes the phosphorylation of PIP2 into PIP3 (PMID: 20622047). PTEN catalyzes the reverse process (PMID: 20622047). We assume that in the absence of PTEN the level of PIP3 follows the level of PI3K.

f_PDK1_pm_=PIP3 or PIP3_2

PIP3 recruits PDK1 to the membrane (PMID: 9895304).

f_mTORC2_pm_=(PIP3 or PIP3_2) and not Everolimus

mTORC2 is stimulated by PIP3 and directed to be co-localized with AKT in the plasma membrane (PMID: 24385483). It is inhibited by Everolimus analogues.

f_AKT_=(PIP3 or PIP3_2) and (PDK1_pm or mTORC2_pm) and (not Ipatasertib or PIP3_2)

PIP3 recruits AKT to the membrane (PMID: 9895304). Membrane-bound PDK1 and mTORC2 phosphorylate AKT at two different sites (PMID: 9094314, PMID: 15718470). We assume that either activation mechanism is sufficient to AKT following its localization directed by PIP3. The drug Ipatasertib inhibits AKT activity; we assume that this inhibition can be overcome by a high level of PIP3. This is consistent with the observation that AKT activity recovers 24 hours after drug treatment in HER2+ cells (PMID: 21215704, PMID 24436048).

f_p21_p27_T_=FOXO3 or not (MYC_2 or MYC)

FOXO3 transcriptionally upregulates p27 (PMID: 10783894). MYC transcriptionally downregulates p21 and p27 (PMID: 15757889, PMID: 11313917). We assume that there is a base level of transcription of p21 or p27 that happens in the absence of MYC activity, thus we combine the positive and negative regulator with the “or not” rule.

f_p21_p27_=(not AKT and not PIM) or p21_p27_T

AKT phosphorylates p21 and inhibits its activity (PMID 11463845). PIM phosphorylates and inhibits the activity of p21 and p27 (PMID: 12431783, PMID:18593906). We assume that a sufficient amount of active (unphosphorylated) p21 or p27 is present if either of two conditions is satisfied: (i) AKT and PIM are absent or (ii) p21 or p27 transcription is upregulated, thus the total and with it the unphosphorylated p21 and/or p27 level increases.

f_cycE_CDK2_T_=E2F or E2F_2 or E2F_3

f_cycE_CDK2_=not p21_p27 and cycE_CDK2_T

E2F transcriptional activators activate the transcription of S phase promoting genes (PMID: 11257102).

p21 and/or p27 inhibit CDK2 (PMID: 10385618)

f_KMT2D_=not AKT

Phosphorylation by AKT attenuates KMT2D’s methyltransferase activity (PMID 28336670)

f_TSC_=not AKT and not SGK1 and not MAPK_2

Both AKT and SGK phosphorylates TSC2, which leads to its destabilization (PMID: 12172553, PMID: 27451907). High ERK activation also inactivates TSC2 (PMID: 17671177).

f_PRAS40_=not AKT and not PIM

Both AKT and PIM phosphorylate PRAS40, which prevents it binding to mTORC1 (PMID: 17386266, PMID: 27604488).

f_mTORC1_=(not TSC or not PRAS40) and not Everolimus

PRAS40 is a component and negative regulator of mTORC1, PMID: 17386266. TSC2 inhibits mTORC1 activation (through converting Rheb to its inactive, GTP-bound form, PMID: 12172553). We assume that mTORC1 is active not targeted by Everolimus and if either inhibitor is inactive.

f_FOXO3_=(not AKT and not SGK1 and not PIM) and not FOXO3_Ub

f_FOXO3_Ub_=MAPK_2

AKT, SGK1 and PIM phosphorylate FOXO3, which leads to its inhibition (PMID: 10102273, PMID: 27451907, PMID:18593906). High ERK activity leads to the ubiquitin-tagging and then proteosomal degradation of FOXO3 (PMID: 18204439). We represent the status of the FOXO3 ubiquitin proteasomal degradation as the FOXO3_Ub node. This node is among the slow nodes (i.e. it is updated with a lower probability).

f_BIM_=(FOXO3 and not MAPK_2) or BIM_T

FOXO3 activates BIM, PMID: 11050388. ERK phosphorylates BIM, which leads to its proteasomal degradation, PMID: 14555991; we assume that this effect happens at high ERK activity only.

f_BAD_= not AKT and not PIM1 and not (MAPK or MAPK_2)

AKT, PIM1 and ERK phosphorylate BAD, which leads to its binding to 14-3-3 proteins and sequestration (PMID: 16226704, PMID: 27604488, PMID: 16226704).

f_MCL1_= translation

MCL-1 is translated in a cap-dependent manner (PMID: 27974663). We assume that this is the main mode of regulation of this anti-apoptotic protein.

f_EIF4F_=mTORC1

mTORC1 (through phosphorylating 4EBP1) allows EIF4E to initiate cap-dependent translation, PMID: 19339977

f_S6K_=mTORC1

mTORC1 activates S6K, PMID: 19339977

f_translation_=EIF4F and S6K

EIF4F initiates cap-dependent translation, PMID: 19339977. S6K activity leads to an increase in mRNA biogenesis and cap-dependent translation, PMID: 19339977.

f_ESR1_=(ER or FOXO3) and not Fulvestrant

f_ESR1_2_=ER and FOXO3 and not Fulvestrant

Cells can have an intrinsic ESR1 level, reflected in the variable ER. Alternatively, FOXO3A can bind to the ESR1 promoter and promote transcription, PMID: 25877889. Fulvestrant is a drug inhibitor of ESR1.

f_FOXA1_= FOXO3

We assume that the pioneer factor FOXA1 is upregulated by FOXO3. This is supported by the existence of a FOXO3 binding site in the promoter region of FOXA1 (based on DECODE by SABiosciences, QIAGEN; http://www.sabiosciences.com/chipqpcrsearch.php?species_id=0&nfactor=n&ninfo=n&ngene=n&B2=Search&src=genecard&factor=Over+200+TF&gene=FOXA1).

f_ER_transcription_=ER and (ESR1 or ESR1_2)

f_ER_transcription_2_=KMT2D and FOXA1 and PBX1 and ESR1_2

The transcriptional regulatory activity of ER is highly enhanced by an open chromatin state (mediated by the histone H3 methyltransferase KMT2D) and the binding of the co-activators FOXA1 and PBX1 (PMID: 28336670)

f_MYC_=ER_transcription

f_MYC_2_=ER_transcription_2

MYC is a key ER-dependent transcriptional target, PMID: 11136970.

f_cyclinD_=MYC

f_cyclinD_2_=MYC_2

MYC activates the transcription of CDK4 and of cyclin D, PMID: 10688915

f_BCL2_=ER_transcription_2 or BCL2_T

In addition to intrinsic expression (incorporated in the source node BCL2_T), BCL2 can be upregulated by ER-dependent transcription (PMID: 21677677, PMID: 20154269, PMID: 11738551).

f_CDK4/6_=not Palbociclib

f_cycD_CDK4/6_=(cyclinD or cyclinD_2) and CDK4/6 fcycD_CDK4/6_2=(cyclinD_2) and CDK4/6

CDK4/6 is present unless inhibited by Palbociclib. CDK4/6 form a complex with the D-type cyclins PMID: 7610482; we assume that the complex has a higher activity when cyclinD has a higher activity.

f_pRb_=(cycD_CDK4/6_2 or cycD_CDK4/6) or cycE_CDK2

f_pRb_2_=(cycD_CDK4/6 and cycE_CDK2) or cycD_CDK4/6_2

f_pRb_3_=cycD_CDK4/6_2 and cycE_CDK2

CDK4/6 hyper-phosphorylates and inhibits Rb, PMID: 11257102. The cyclinE/CDK 2 kinase co-operates with cyclin D/CDK4/6 to phosphorylate Rb (PMID: 10323868). The highest level of hyper-phosphorylation is when both are at their most active levels.

f_E2F_=pRb

f_E2F_2_=pRb_2

_fE2F_3_=pRb_3 or (pRb_2 and E2F_3)

Unphosphorylated Rb binds to the E2F family of transcription factors, preventing it from interacting with the cell’s transcription machinery (PMID: 11257102). Thus the (hyper)phosphorylated forms of Rb activate E2F. High E2F activity can be sustained by self-regulation, as E2F family members can activate their own transcription (PMID: 21677677, PMID: 18364697).

f_proliferation_=translation or E2F or E2F_2 or E2F_3

f_proliferation_2_=translation or E2F_2 or E2F_3

f_proliferation_3_=(translation and E2F_2) or E2F_3

f_proliferation_4_=translation and E2F_3

The output node proliferation focuses on two aspects of cell proliferation: cell cycle transition from G1 to S (reflected in the level of E2F), and protein translation. The lowest level of proliferation propensity is assumed in the presence of either of these; the highest corresponds to the simultaneous presence of protein translation and the highest activity level of E2F.

f_apoptosis_=(BIM and not (MCL1 and BCL2)) or (BIM and BAD) or (BAD and not (MCL1 and BCL2)) or apoptosis

f_apoptosis_2_=(BIM and BAD and not (MCL1 and BCL2)) or apoptosis_2

f_apoptosis_3_=((BIM and BAD and not (MCL1 or BCL2))) or apoptosis_3

The anti-apoptotic proteins BCL2 and MCL1 inhibit apoptosis effectors, while the apoptosis sensitizer BAD and the apoptosis activator BIM activate apoptosis effectors (PMID: 25723171, PMID: 25895919). We assume the lowest level of apoptosis propensity when the activation of a pro-apoptotic protein is coupled with the inactivity of one of the anti-apoptotic proteins, or alternatively when both types of pro-apoptotic proteins are active. The highest level of apoptosis propensity corresponds to the case when both types of pro-apoptotic proteins are active and both anti-apoptotic proteins are inactive. We assume that the apoptosis propensity cannot decrease (i.e. the apoptosis priming process is irreversible).

#### Initial or externally controlled states

The source nodes that correspond to drug treatment (Alpelisib, Fulvestrant, Neratinib, Palbociclib, Everolimus, Trametinib, Ipatasertib) are switched on according to the modeled treatment scenario.

Within the 13 source nodes, IGFR_T and PBX1 are assumed to be ON, assuming an above-threshold intrinsic level. ER is assumed to be ON, HER2 and HER3 are OFF, to reflect an ER+ /HER2-cell. CDK4/6, PTEN, SGK1_T, PIM, PDK1, and mTORC2 (which act as resistance mechanisms to PI3K inhibitors) are OFF in the default case and assumed to be ON when simulating the relevant resistance scenario. The apoptotic protein transcripts BIM_T and BCL2_T can be either ON or OFF.

The long-term behaviors (attractors) of the model under the selected source node states are obtained using stable motif analysis (PMID: 25849586), which results in 8 steady state attractors: 6 cancerous steady states with high survivability (Apoptosis = 0, Proliferation = 3 or 4) and 2 cancerous steady states with lower survivability (Apoptosis = 1, Proliferation = 3 or 4). The initial state during simulations corresponds to a mix of two of the high survivability cancerous states (see Fig. 4A): PI3K and PIP3, AKT, RAS and MAPK is 1. FOXO3, PRAS40 and TSC are OFF. mTORC1 and the rest of the mTORC1 pathway are ON. ER_transcription, MYC and cyclinD, CDK4/6, and cycD_CDK4/6 are at level 1. p21/p27 and p21/p27_T are OFF, cycE/CDK2 and cycE/CDK2_T are OFF, Rb is phosphorylated, E2F is at level 2, and proliferation is at level 3. BAD is OFF, MCL1 is ON, BIM and BCL2 are simultaneously ON in 1/3 of the simulations (and OFF in the rest), apoptosis is at level 0.

#### Update probability of the nodes

The source nodes that correspond to drug treatment (Alpelisib, Fulvestrant, Neratinib, Palbociclib, Everolimus, Trametinib, Ipatasertib) are switched on according to the modeled treatment scenario.

We categorize nodes into fast or slow depending on whether the node is activated by a (fast) signaling event or a (slow) transcriptional/translational event, and set the update probability of fast nodes to be 5 times higher than that of slow nodes. To aid model expansion, this categorization includes source nodes whose regulatory function is f_X_=X in the current version of the model.

The fast nodes are: AKT, BAD, cycE_CDK2, EIF4F, FOXO3, HER2/HER3, KMT2D, MAPK, mTORC1, mTORC2_pm, p21_p27, PDK1_pm, PI3K, PIM, PIP3, PRAS40, Proliferation, RAS, S6K, SGK1, Translation, TSC.

The slow nodes are: Apoptosis, BCL2, BCL2_T, BIM, BIM_T, CDK4/6, cycD_CDK4/6, cycE_CDK2_T, cyclinD, E2F, ER, ER_transcription, ESR1, FOXA1, FOXO3_Ub, HER2, HER3, HER3_T, IGF1R, IGF1R_T, MCL1, mTORC2, MYC, p21_p27_T, PBX1, PDK1, pRb, PTEN, SGK1_T.

#### Key experimental and clinical outcomes reproduced by the model

The model recapitulates a variety of observations in ER+ and HER2+ breast cancer. A selected group of key observation are given below:

Drug inhibition of MEK in HER2+ breast cancer cells leads to increased HER2/HER3 heterodimer formation and higher PI3K activation (PMID: 22552284).

High HER3 expression induces resistance to PI3K inhibitors, which is overcome by HER3 blockade, in HER2-amplified and/or PIK3CA-mutant breast cancer cell lines and brain metastases of mouse xenografts (PMID: 28539475).

Inhibition of AKT (directly by AKT inhibitors or indirectly by mTOR or PI3K inhibitors) induces the activity of the transcription factor FOXO3, which upregulates a shared set of RTKs, including HER3, IGF1R, in HER2+ breast cancer cell lines and mouse xenografts tumors (PMID: 21215704, 22140653, 21368164).

High PIM expression is a resistance mechanism to PI3K inhibitors in ER+ (PIM1/2/3) and HER2+ (PIM2) breast cancer cell lines. High PIM1/3 expression is observed in biopsies of ER+ human tumors treated with PI3K inhibitors (PMID: 27604488, 28561737).

High PDK1/SGK1 expression is a resistance mechanism to PI3K inhibitors in HER2+ breast cancer cell lines and mouse xenografts tumors. High SGK1 expression and activity in breast cancer tumor samples causes intrinsic resistance to PI3K inhibitors (PMID: 27451907).

High PDK1 and AKT2 expression are putative resistance mechanisms to PI3K inhibitors; they are observed in biopsies of ER+ human tumors treated with PI3K inhibitors (PMID: 27604488).

Inhibition of PI3K induces a rapid downregulation of MAPK signaling and induction of apoptosis in ER+ and HER2+ breast cancer cell lines and mouse xenograft tumors. In HER2+ breast cancer cell lines, MAPK activity is reactivated following the induction of RTKs (PMID: 24436048, 24327733, 25544637).

Inhibition of PI3K in ER+ breast cancer cell lines induces the transcription factor activity of FOXO3, which binds the promoters of ESR1 and HER3, and upregulates their expression. The upregulation of ESR1 expression in response to PI3K inhibitors has also been observed in ER+ mouse xenograft tumors and ER+ human breast cancer tumor biopsies (PMID: 25877889, 28539475).

## Footnotes

1 Here we use the term “inhibitor” to refer to drugs that target and inhibit an element regardless of the specific mechanism of action.

